# Short-term caloric restriction or resveratrol supplementation alters large-scale brain network connectivity in male and female rats

**DOI:** 10.1101/2024.10.02.616243

**Authors:** Judith R.A. van Rooij, Monica van den Berg, Tamara Vasilkovska, Johan Van Audekerke, Lauren Kosten, Daniele Bertoglio, Mohit H. Adhikari, Marleen Verhoye

## Abstract

**Introduction:** Dietary interventions such as caloric restriction (CR) exert positive effects on brain health. Unfortunately, poor compliance hinders the success of this approach. A proposed alternative is resveratrol (Rsv), a CR-mimetic known to promote brain health. Direct comparison between the effects of Rsv and CR on brain health is lacking, with limited knowledge on their sex-specific effects. Therefore, we aimed to compare and unravel the sex-specific impact of these dietary interventions on spontaneous brain activity.

**Methods:** Here, we used resting-state fMRI to investigate functional connectivity (FC) changes in five prominent resting-state brain networks (RSNs) in healthy four month old male and female F344 rats supplemented to either 40% CR or daily Rsv supplementation (10 mg/kg, oral) for the duration of one month.

**Results:** Our results demonstrated a decreased body weight (BW) in CR rats, as well as an increase in body weight in male Rsv supplemented rats, compared to female Rsv supplemented rats, whereas this difference between sexes was not observed in the control or CR groups. Furthermore, we found that both CR or Rsv supplementation induce a female-specific decrease of FC between the subcortical network and hippocampal network, and between the subcortical network and lateral cortical network. Moreover, Rsv supplementation lowered FC within the hippocampal network and between the hippocampal and the default mode like network, the lateral cortical network and the sensory network – an effect not observed for the CR rats. Finally, we observed an overall lower FC in male rats compared to females, irrespective of dietary intervention, within the subcortical network and between the subcortical, the sensory and default mode like network.

**Discussion:** Our findings reveal that both CR and Rsv induce a similar female-specific decrease of FC in RSNs associated with memory and emotion, all the while CR and Rsv induce dissimilar changes in body weight and other within- and between RSN FC measures. Altogether, this study provides insight into the effects and comparability of short-term CR and Rsv supplementation on brain connectivity within and between RSNs in both male and female F344 rats, providing a FC reference for future research of dietary effects.

## 1. Introduction

Due to the constant exponential growth of the elderly population world-wide, the search for preventative strategies to preserve cognitive function and promote healthy aging have become increasingly important (1). Lifestyle modifications, including regular exercise and dietary adjustments, are known to play a crucial role in this endeavor. One proposed dietary adjustment is caloric restriction (CR) (2), defined as a significant reduction of energy intake without the risk of malnutrition. Through a variety of pathways, short- and long-term CR has been shown to positively influence brain health by reducing neuroinflammation (3) and oxidative stress (4), while simultaneously enhancing neurovascular function (5). In animal models of aging and age-related neurodegenerative diseases, CR has been shown to prevent neuronal degradation (6, 7), enhance neurogenesis in the hippocampus (8), as well as prevent against the age-related decline in motor coordination (9) and learning (7). These findings collectively highlight the potential of CR as a promising approach to maintain and improve brain health.

Despite the proven positive effect of CR on brain health, poor compliance to drastic, life-long CR hinders its success as a therapeutic approach (10). In light of this challenge, CR-mimetics have emerged as a potential alternative strategy. CR-mimetics aim to activate the same pathways and mechanisms and are therefore hypothesized to mimic the beneficial effects of CR on brain health without the life-long commitment to reduced caloric intake (11, 12). One of the most extensively studied CR-mimetics is resveratrol (Rsv), a polyphenol and phytoalexin, present in high concentrations in, for example, blue berries and red grapes. Rsv has shown to exert positive effects on neuroinflammation (13) and oxidative stress (14). The hypothesis of Rsv serving a similar function as CR is further solidified through reported findings from studies in animal models of aging and age-related neurodegenerative diseases (15). Here, Rsv is reported to preserve cognitive function (16) and neurovascular coupling (17) in aging mice, and exert neuroprotective effects in neurodegenerative diseases such as Alzheimer’s (18) and Parkinson’s Disease (19) through similar pathways as reported for CR. These findings further support the hypothesis that Rsv may serve as a viable alternative to CR in promoting brain health and potentially delaying the onset of aging and age-related neurodegenerative diseases.

Resting-state functional MRI (rsfMRI), a non-invasive neuroimaging method, has proven itself as a powerful tool to assess brain function (20). With rsfMRI, one measures the blood-oxygenation level dependent (BOLD)-contrast fluctuations, indirectly reflecting fluctuations in neuronal activity via neurovascular coupling while the brain is at rest. The correlation between the BOLD signal timeseries of brain regions is expressed as functional connectivity (FC), where regions that exhibit highly correlated BOLD signals form the resting-state networks (RSNs) (21). The most prominent RSN in humans is the default mode network (DMN), typically anticorrelated to the frontoparietal network (FPN) (22). Changes in FC as a result of CR and Rsv have been reported in regions belonging to the hippocampal network (Hipp), FPN and the DMN (23-26), emphasizing the potential of rsfMRI to assess changes in brain function as a result of dietary interventions.

While these findings confirm that CR or Rsv supplementation alters brain connectivity, there is a lack of research directly comparing the effects of these dietary interventions. Moreover, even though both CR and Rsv supplementation have been hypothesized to exert effects on brain function in a sex-dependent manner (27, 28), research on the possible sex-specific effects that both dietary interventions can exert on brain connectivity, is sparce. Clinical trails assessing dietary effects are subject to challenge due to low patient adherence, high susceptibility to confounding variables, high patient dropout rates and limited follow-up periods (29). The use of animal models can overcome these challenges, aiding to provide critical insight into the sex-specific effects of CR or Rsv on brain connectivity. RsfMRI is well established in rodents, showing similar findings compared to humans regarding RSNs. Rodent analogues of the human DMN and FPN, along with other major RSNs, have been identified as the default mode-like network (DMLN), the lateral cortical network (LCN), the hippocampal (Hipp), the sensory (Sens) and subcortical (SubC) network (30). In this study, we hypothesized that CR or Rsv supplementation induces changes in connectivity within- and between these RSNs in a sex-specific manner. Therefore, we aimed to characterize and compare the sex-specific effects of short-term CR and Rsv supplementation on brain connectivity, using rsfMRI to assess FC changes within- and between the five prominent rodent RSNs as a result of dietary intervention.

## 2. Materials and methods

### 2.1. Animals

F344 rats (RRID: RGD_60994, Charles River, Italy) used within this study were bred in-house. A total of 42 male and female F344 rats (n=21/sex) were single-housed at three months of age for the duration of the experiment, and kept under controlled environmental conditions (12-h light/dark cycle, (22 ± 2)°C, 40-60% humidity). Water was provided at libitum. All procedures were in accordance with the guidelines approved by the European Ethics Committee (decree 2010/63/EU), and were approved by the Committee on Animal Care and Use at the University of Antwerp, Belgium (ECD: 2021-59).

### 2.2. Dietary treatment paradigm

Three-month-old rats were randomly divided into three groups (Figure 1), receiving dietary intervention and Rsv supplementation for the duration of four weeks, until the age of four months. The first group served as controls. The second group received daily Rsv (10 mg/kg, oral, High Potency Trans-Resveratrol 600, Doctor’s Best®, California, USA) supplementation. The third group was exposed to a gradual weekly decrease in caloric uptake by 40%. To allow adaptation to restricted feeding, the animals were fed one week with 15% CR, a second week with 27.5% CR and 40% CR thereafter. Food fortified with vitamins and minerals was provided to avoid malnutrition (Rat/Mouse Fortified, Ssniff®, Germany). CR of 15%, 27.5% and 40% was calculated based on averaged food intake of Ctrl (male/female n = 3/3) and Rsv (male/female n = 3/2) rats, adjusted for age and sex. Control and Rsv supplemented rats were provided standard chow (Ssniff®, Germany) ad libitum. Rsv solubilized in 99.8% ethanol (Thermo Fisher Scientific), was applied to dried apple chips (JR Farm, Germany) and supplemented daily to the rats in the Rsv group. To minimize the difference between conditions, all CR and control rats were supplemented with a daily dose of dried apple chips with vehicle only (99.8% ethanol). This volume of ethanol was determined through averaging the daily Rsv volumes, adjusted for age and sex.

**Figure 1:**
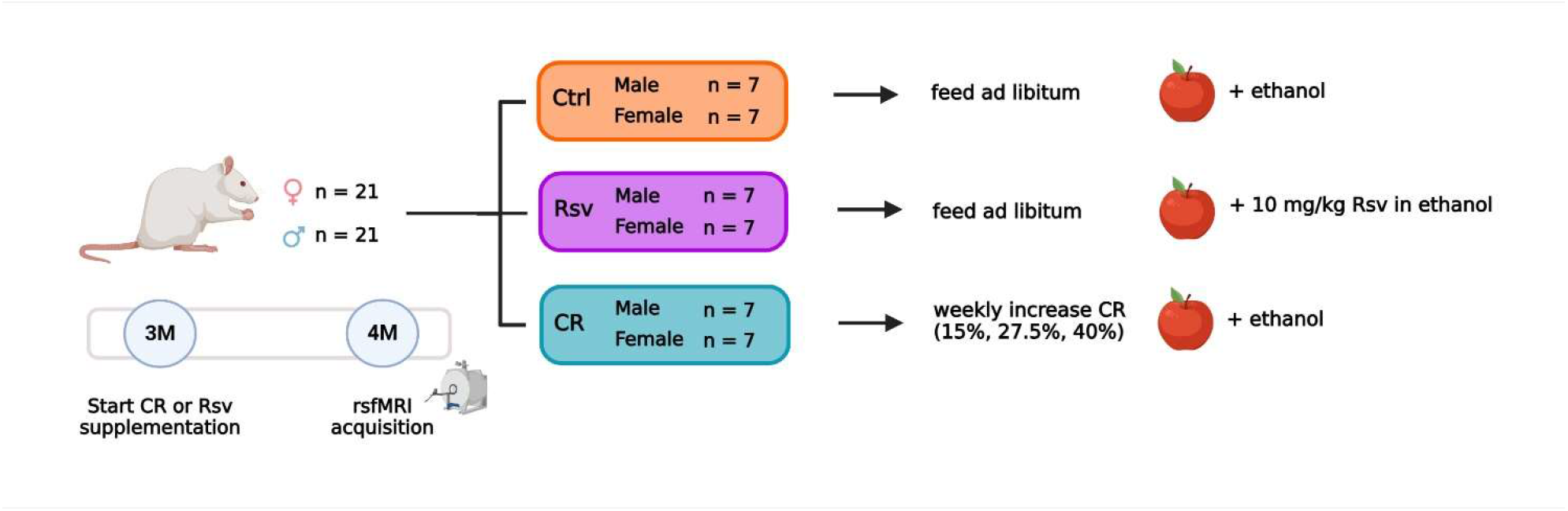
Overview of the experimental dietary treatment paradigm and study design. Number of animals per sex and group are given. Dietary intervention and supplementation of the apple chips starts at three months of age (3M). RsfMRI data are acquired at four months of age (4M). The apple represents the supplementation of Rsv or its vehicle using dried apple chips. Created with Biorender.com

Subjects were weighed prior to the start of the dietary intervention and each week afterwards for the duration of the experiment. The percentage body weight (BW) change respective to the starting weight was calculated for each subject.

### 2.3. RsfMRI acquisition

RsfMRI data of the rats were acquired at four months of age, after 1 month of CR or Rsv supplementation. Rats were anaesthetized using an isoflurane-medetomidine protocol: an established anesthesia protocol for rsfMRI in rodents (31-34). Anesthesia was induced using 5% isoflurane (IsoFlo®, Zoetis, US), administered with a gaseous mixture of 200 ml/min O_2_ and 400 ml/min N_2_. For further animal handling and positioning, the isoflurane level was reduced to 3%. After positioning, a subcutaneous bolus injection of medetomidine (0.05 mg/kg, Domitor®, Vetoquinol, France) was administered. Continuous subcutaneous infusion of medetomidine (0.1 mg/kg/h) started 15 minutes after bolus administration, with the isoflurane level being lowered to 0.4%, to be maintained throughout the whole MRI session. Physiological parameters (breathing rate, heart rate, O2 saturation and body temperature) of the animal were closely monitored during the entire procedure (MR-compatible Small Animal Monitoring, Gating, and heating system, SA Instruments). Body temperature was maintained at 37 (± 0.5) °C using a feedback-controlled warm air circuitry (MR-compatible Small Animal Heating System, SA Instruments Inc., USA).

MRI data were acquired on a 7T Pharmascan MR scanner (Bruker®, Germany) with a volume resonator coil for radiofrequency excitation and a 2×2 channel receiver head radiofrequency coil for signal detection. To ensure uniform slice positioning between subjects, multi-slice T2-weighted (T2w) TurboRARE images were acquired in three directions (echo time (TE): 33 ms, repetition time (TR): 1800 ms, RARE factor: 8, field of view (FOV): (35 × 35) mm^2^, matrix: [256 x 256]). Whole-brain rsfMRI data were acquired 40 minutes after the bolus of medetomidine using a single shot gradient echo echo-planar imaging (EPI) sequence (TE: 18 ms, TR: 600 ms, FOV (30 x 30) mm^2^, matrix [96 x 96], 12 coronal slices of 1 mm, slice gap: 0.1 mm, 1000 repetitions), for a total of 10 minutes. An anatomical 3D image was acquired for registration purposes, with a T2w-TurboRARE sequence (TR: 1800 ms, TE: 36 ms, RARE factor: 16, FOV: (35 x 35 x 16) mm^3^, acquisition matrix: [256 x 256 x 32], reconstruction matrix: [256 x 256 x 64]). At the end of the scan session, a subcutaneous injection of 0.1 mg/kg atipamezole (Antisedan®, Pfizer, Germany) was administered to counteract the effects of the medetomidine anesthesia and the animals were allowed to recover under a heating lamp. All animals recovered within 15-20 minutes after the end of the scan session.

### 2.4. rsfMRI image preprocessing

All preprocessing steps were performed with MATLAB R2020a (Mathworks, Natick, MA) and ANTs (Advanced Normalization Tools). The rsfMRI data were padded using an in-house MATLAB script. Debiasing, realignment, normalization, co-registration and smoothing of the data was performed using SPM 12 software (Statistical Parametric Mapping). First, subject-specific 3Ds were debiased after which a study specific 3D-template was created from a subset of animals (male/female: Ctrl n = 2/2, Rsv n = 2/2, CR n = 2/3) in ANTs. All individual 3Ds were normalized to the study-specific 3D template using a global 12-parameter affine transformation followed by a non-linear deformation protocol. The rsfMRI EPI images were realigned to the first EPI image using a 6-parameter rigid body spatial transformation estimated using a least-squares approach. RsfMRI data were co-registered to the animal’s respective 3D image using a global 12-parameter affine transformation with mutual information used as similarity metric and normalized to the study-specific template using the combined transformation parameters. RsfMRI data were smoothed in-plane using a Gaussian kernel with full width at half maximum of twice the voxel size and filtered (0.01 – 0.2 Hz) with a Butterworth band-pass filter. Finally, quadratic detrending was performed on the filtered images. During the entire process, a total of five subjects (Ctrl (male/female n = 2/1), CR (male n = 2)) were removed from further analysis due to poor quality of the data.

### 2.5. Functional connectivity (FC) analysis

Region of interest (ROI)-based FC analysis was performed on the preprocessed rsfMRI data. A neuroanatomical atlas comprised of 71 anatomical parcels (Fischer 344, https://www.nearlab.xyz/fischer344atlas), was warped onto the study-specific template and down-sampled (ANTs) to match the EPI space. Out of the 71 parcels, 43 unilateral grey matter ROIs (for both the left (L) and right (R) hemisphere for each region) were selected, excluding regions sensitive to susceptibility artefacts, transient effects and small size. These selected ROIs represent the five prominent rodent RSNs: the default mode like network (DMLN), the hippocampal network (Hipp), the sensory network (Sens), the lateral cortical network (LCN) and the subcortical network (SubC) (Supplementary Table 1). For each subject, Pearson correlation coefficients between the ROI-averaged BOLD signal timeseries of each pair of ROIs were calculated and Fisher z-transformed yielding subject-wise 43×43 FC matrices. FC within each network was calculated by averaging across FC values between all ROIs belonging to the network. Similarly, between-network FC was calculated by averaging across FC values between all pairs of ROIs belonging to both networks.

### 2.6. Statistics

The ROI-based and network-based FC matrices were subjected to an one-sample t-test (FDR corrected, Benjamin-Hochberg procedure, p < 0.05), within group (per sex and treatment). All data was tested for normality using the Shapiro-Wilk test determining the goodness of a normal fit. All data was normally distributed. The BW at week 0, the percentile change in BW at 4 weeks of treatment and the outcomes of the network-based FC were analysed using a two-way ANOVA (treatment, sex, treatment*sex). In case of a significant treatment*sex interaction, post-hoc tests were performed using Student’s t-test with FDR correction (Benjamin-Hochberg procedure, p < 0.05). When no significant treatment*sex interaction was present, the interaction was removed and the model was recalculated using only the main effects (treatment and sex). In case of a significant treatment effect, post-hoc tests were performed on all groups using a Tukey HSD test. Outlier detection was performed using a principal component analysis, per sex and treatment. Subjects with a Hotelling T2 statistics index higher than the 95% confidence interval were marked as outliers. Subjects with more than 8 out of 15 within- and between network FC values marked as outliers were excluded. Statistical analyses were performed using JMP Pro 17 (SAS Institute Inc.) and MATLAB R2020a (Mathworks, Natick, MA). Graphical representation of the data was created using GraphPad Prism (version 9.4.1. for Windows, GraphPad Software, San Diego, California USA) and Adobe Illustrator (Adobe Inc.).

## 3. Results

### 3.1. Caloric restriction reduces body weight in both male and female rats

In order to test the randomness of the treatment groups prior to dietary intervention, we tested for the effects of treatment and sex on body weight at week 0. No significant effects were found for treatment (p = 0.4324) or the interaction treatment*sex (p = 0.6471), but, as expected, we did find a significant sex effect (p < 0.0001) on BW, where males had a significantly higher weight compared to females (Figure 2A).

**Figure 2:**
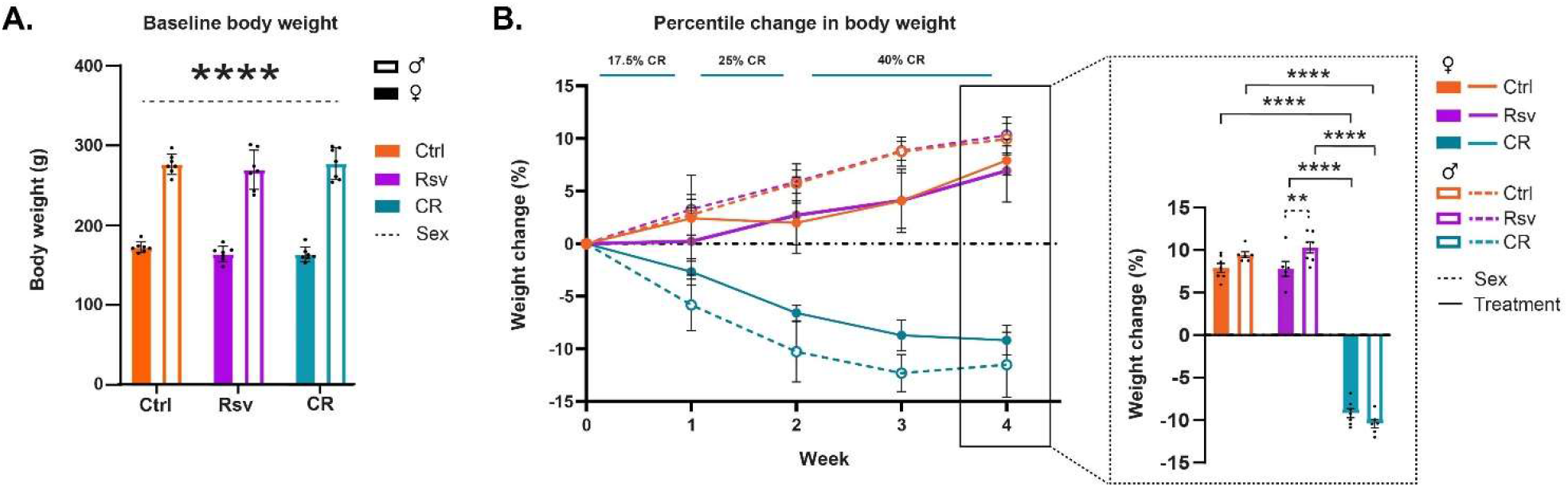
Effects of dietary intervention on body weight (BW). **(A)** Mean ± SD BW in grams (g) at baseline, prior to the start of the dietary interventions (week 0). The colors of the bars are indicative of the group each subject was assigned to (control (Ctrl), resveratrol (Rsv), caloric restricted (CR)). Solid bars represent females and open bars represent the males. Dots represent individual subject data points. **(B)** Percentage change in BW (mean **±** SD) with respect to baseline, over the course of the 4 week treatment, Ctrl, Rsv and CR group. Bar graph (insert) shows the percentage change in BW (mean **±** SD) at week 4. Asterisks indicate the levels of statistical significance: **p<0.01, ****p<0.0001.

Next, we aimed to determine the effect of short-term dietary intervention on BW, by evaluating the percentile change in BW over the course of treatment, from the start of the intervention (Figure 2B). Over the course of the intervention, we observed a positive percentage change of BW in both Ctrl and Rsv rats indicative of weight gain, while observing a negative percentage BW change in the CR rats, indicative of weight loss. Statistical analysis of the percent change in BW at week 4 demonstrated a significant treatment*sex interaction effect (p = 0.031). Post-hoc analysis revealed that after four weeks the percentage BW change was significantly different in CR female rats ((*mean ± SD)* %), (-9.18 ± 1.41)%) when compared to female Ctrl ((6.94 ± 2.98)%, p < 0.0001) and female Rsv (7.92 ± 1.44)%, p < 0.0001) rats. Similarly, we observed a significant difference in male CR rats (-11.51 ± 3.73)% compared to male Ctrl ((9.95 ± 1.67)%, p < 0.0001) and male Rsv ((10.33 ± 1.71)%, p < 0.0001) rats. Moreover, male Rsv supplemented rats had a higher percentage increase in BW compared to the female Rsv supplemented rats (p = 0.009). This sex-difference was not observed in the Ctrl (p = 0.1084) or CR (p = 0.0739) groups.

### 3.2. Dietary interventions alter ROI- and network-based resting-state FC

Next, we investigated the effect of dietary intervention on brain connectivity. To evaluate these effects on FC between ROIs and within and between networks, we first calculated average ROI-based FC matrices between the 43 predefined ROIs, per sex and treatment (Figure 3A), providing insight into the distribution of FC across the groups. Our analysis revealed that within group, FC of regions associated with the DMLN, Hipp, Sens and LCN were statistically significant and thus robustly represented in each of the groups. ROI-based FC, mainly within the SubC (caudate putamen (CPu), medial septum (MS), thalamus (Thal), hypothalamus (Hyp) and nucleus Accumbens (nAcc)), were often not statistically significant in the males, irrespective of treatment. The total number of significant connections was higher in females when compared to males in each group. An additional graphical representation of the ROI-based FC matrices was generated (Supplementary Figure 1), showing strong inter-hemispheric FC within the DMLN and LCN, and intra-hemispheric FC between DMLN, Hipp and Sens. These observations were primarily noted for the Ctrl and CR groups and to a lesser extent in the Rsv supplemented group.

**Figure 3:**
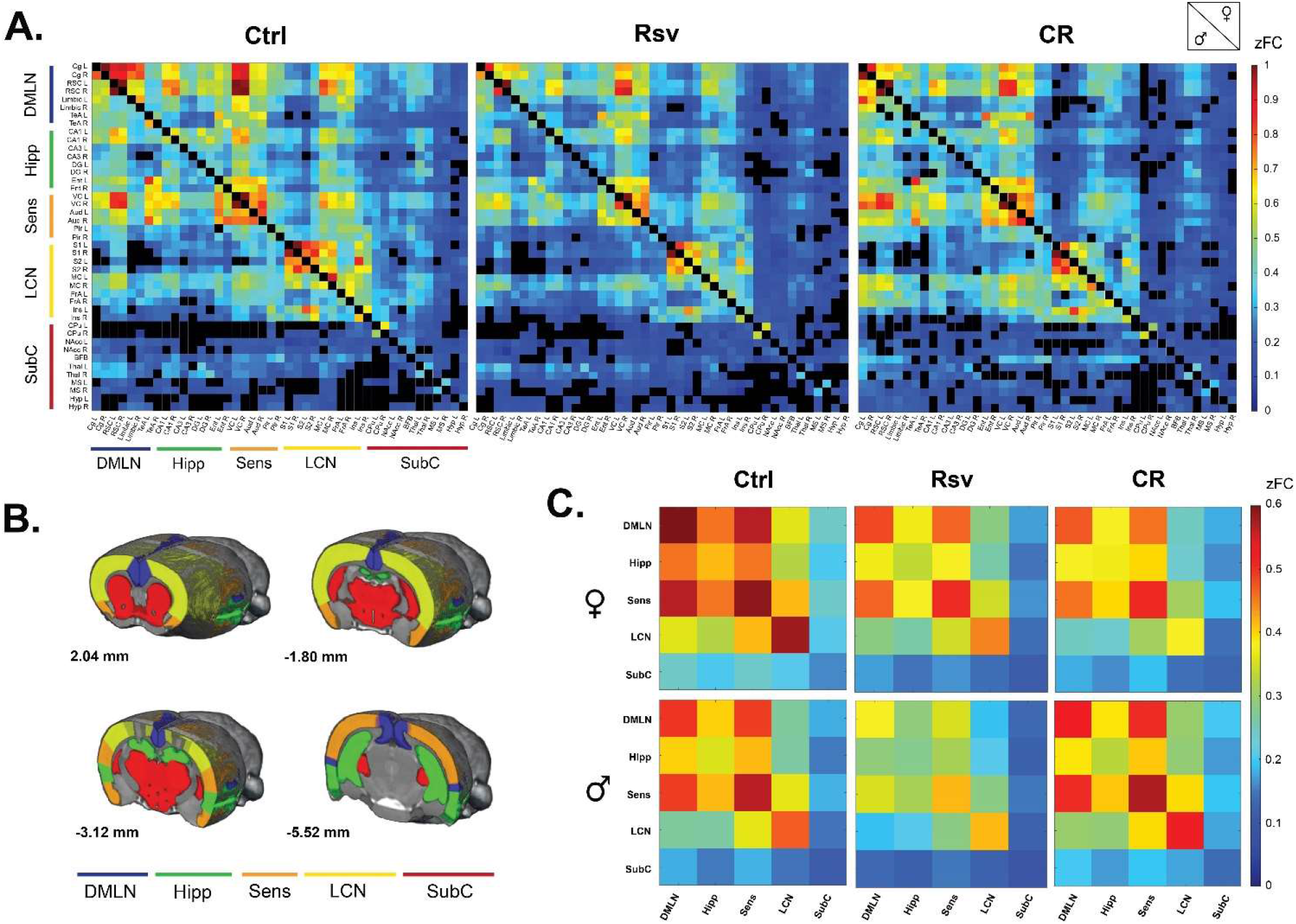
FC in male and female Ctrl, Rsv supplemented and CR rats. **(A)** ROI-based FC matrices displaying the mean FC between ROI pairs for males (lower half) and females (upper half) per treatment group (Ctrl = control, CR = caloric restriction, Rsv = Resveratrol, zFC = z-scored functional connectivity). The 43 regions of interest with left (L) and right (R) hemispheric location are plotted on the x and y axes (Supplementary Table 1). Non-significant connections (p> 0.05, one-sample t-test, FDR corrected per group) are blacked out. **(B)** Anatomical representation of the RSNs used in the network-based FC analysis. Annotated per image are the anatomical Bregma depths. **(C.)** Mean network-based FC in females (top row) and males (bottom row) per treatment group. Colors indicate the strength of FC. Off-diagonal connections represent between-network FC, whereas the diagonals represent within network FC: DMLN (default mode-like network), Hipp (hippocampal network), Sens (sensory network), LCN (lateral cortical network), SubC (subcortical network). All within and between network FC values were significantly different from 0 as determined with a one-sample t-test (FDR corrected, Benjamin-Hochberg procedure, p<0.05).

Next, we grouped the ROIs based on their anatomical location and known implication into five RSNs (Figure 3B) and calculated, similar to the ROI-based FC matrices, average network-based FC matrices per sex and treatment (Figure 3C). Despite the earlier mentioned lack of significance in regions associated with the SubC, all between and within network FC measures were significantly higher than zero within each group (p < 0.05, one-sample t-test, FDR corrected (Supplementary Table 2 and 3)).

To visualize the effects of dietary intervention on ROI-based and network-based FC, we calculated the mean difference between the treatment groups per sex, highlighting the global changes in FC as a result of dietary intervention (Figure 4A and 4B).

**Figure 4:**
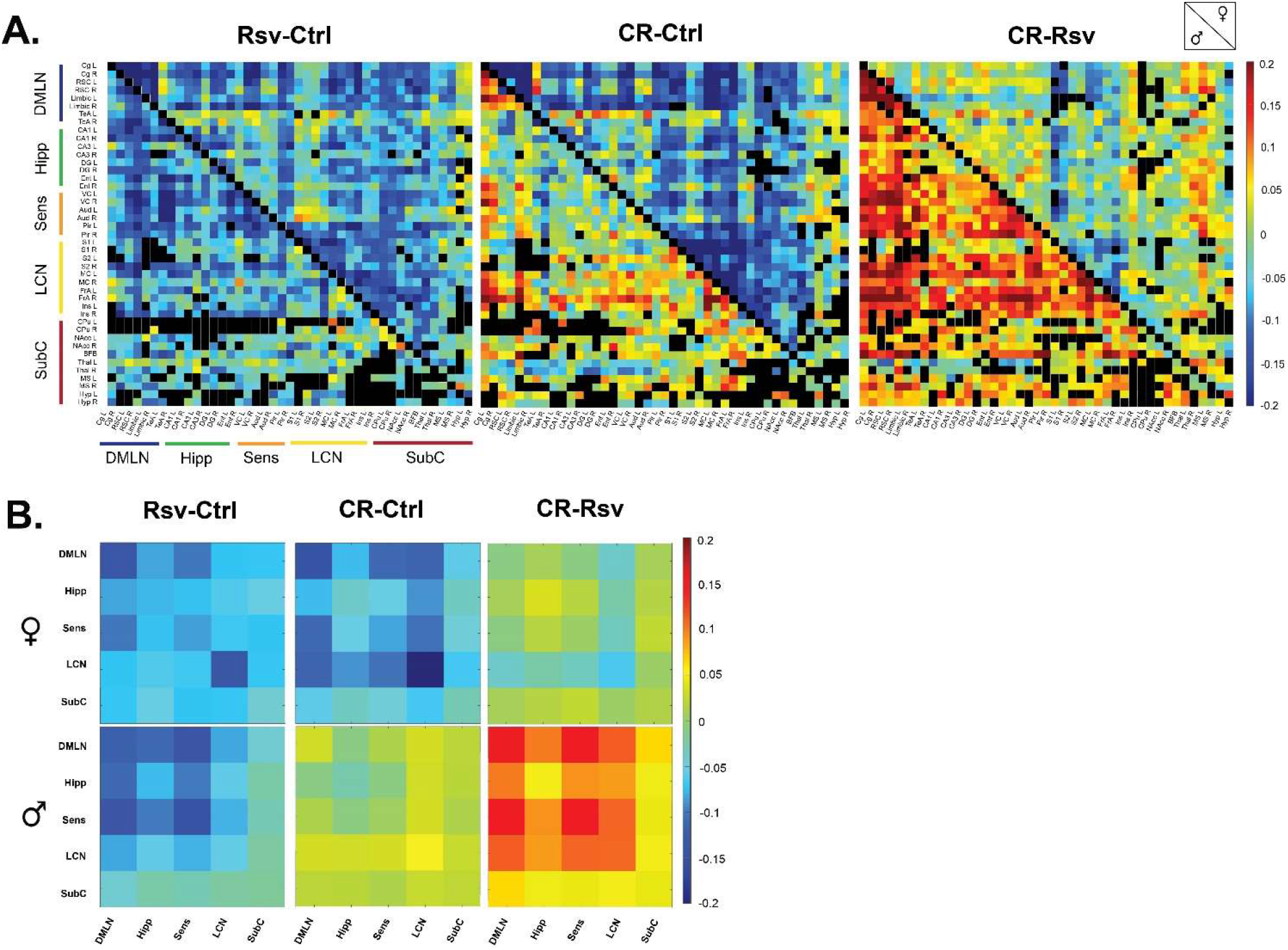
Mean difference in FC in male and female Ctrl, Rsv supplemented and CR rats. (A) ROI-based FC matrices displaying the mean difference in FC between ROI pairs for males (lower half) and females (upper half) between the treatment groups (Ctrl = control, CR = caloric restriction, Rsv = Resveratrol). Non-significant connections (calculated on the mean FC, p>0.05, one-sample t-test, FDR corrected per group) are blacked out. (B.) Mean difference in network-based FC in females (top row) and males (bottom row) between treatment groups. In both panel A and B, positive (red-colour) and negative (blue-colour) values indicate, respectively, higher and lower connectivity in the first group relative to the second group. Diagonal and off-diagonal elements represent differences in within- and between-network FC respectively: DMLN (default mode-like network), Hipp (hippocampal network), Sens (sensory network), LCN (lateral cortical network), SubC (subcortical network).

**Figure 5:**
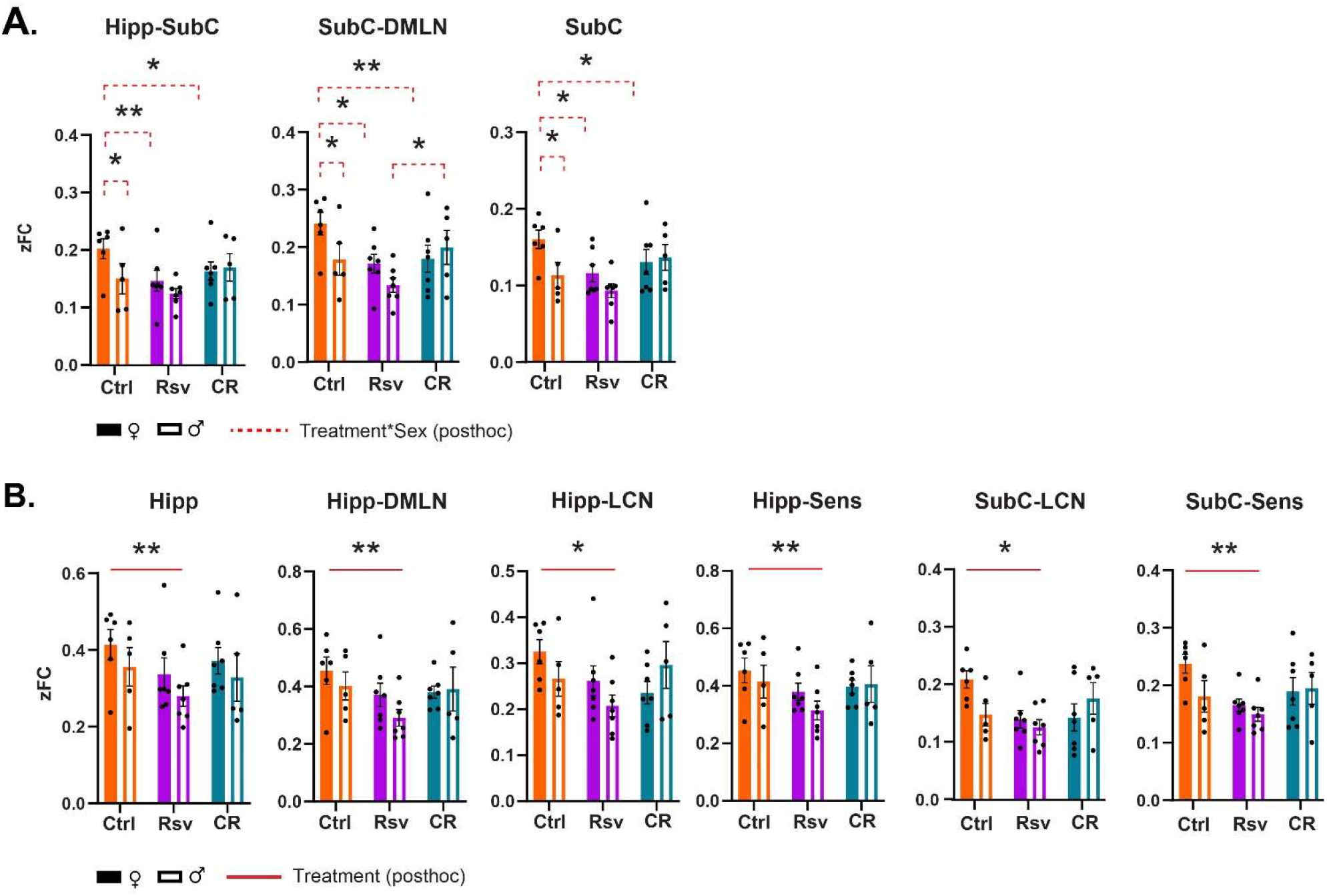
RSN connectivity alterations in male and female Ctrl, Rsv supplemented and CR rats. Network-based FC demonstrating significant interaction **(A)** or treatment **(B)** effects. Bar graphs show the group-level FC within (e.g. Hipp) or between (e.g. Hipp-Sens) FC. Solid bars represent females and open bars representing the males. Group means ± SEM are presented together with the individual subject data points (dots). Significant interaction effects of treatment*sex (FDR corrected, Benjamin-Hochberg procedure, p<0.05) are noted by the red dotted line. Significant treatment effects (post-hoc Tukey HSD p<0.05) are annotated by the red solid line. Asterisks indicate the levels of statistical significance: *p<0.05 **p<0.01, ***p<0.001, ****p<0.0001.

### 3.3. Hippocampal and subcortical within- and between network FC is different between sexes and altered by dietary intervention

To further investigate if FC is altered due to dietary intervention within and between RSNs, data was subjected to statistical analysis (two-way ANOVA). We observed a significant interaction effect of treatment*sex in the Hipp-SubC (p = 0.0250), SubC-DMLN (p = 0.0116) and SubC (p = 0.0215) (Figure A). Hipp-SubC FC was higher in female Ctrl rats when compared to male Ctrl rats (p = 0.0229), female Rsv rats (p = 0.0038) and female CR rats (p = 0.019). Similarly, in the SubC-DMLN, FC was higher in female Ctrl rats compared to male Ctrl rats (p = 0.0364), female Rsv rats (p = 0.0364) and female CR rats (p = 0.0083) and in addition we observe a difference between male CR and male Rsv rats (p = 0.0495). Finally, we also observe this interaction effect in the SubC, where FC was higher in female Ctrl rats compared to male Ctrl rats (p = 0.0225), female Rsv rats (p = 0.0225) and female CR rats (p = 0.0225).

Moreover, we found a significant treatment effect demonstrating lower FC in the Rsv group, compared to the Ctrl group in the Hipp network (p = 0.0036), Hipp-DMLN (p = 0.0030), Hipp-LCN (p = 0.0474), Hipp-Sens (p = 0.0055), SubC-LCN (p = 0.0339) and SubC-Sens (p = 0.0076). Post-hoc analysis (Figure B) consistently revealed a lower FC in the Rsv group, when compared to the Ctrl group (Hipp network (p = 0.0025), Hipp-DMLN (p = 0.0020), Hipp-LCN (p = 0.0375), Hipp-Sens (p = 0.0043), SubC-LCN (p = 0.0266) and SubC-Sens (p = 0.0062)) - an effect we did not observe in the CR group (Supplementary Table 4-6).

## 4. Discussion

To our knowledge, this is the first study to characterize and compare RSN FC in both male and female F344 rats subjected to short-term CR or Rsv supplementation, comparing not only the effects of CR and Rsv on RSN FC, but also highlighting their sex-specific effects. Our results demonstrated decreased BW in CR rats, as well as increased in BW in male Rsv supplemented rats, compared to female Rsv supplemented rats, whereas this difference between sexes was not observed in the Ctrl or CR groups. Furthermore, we found that both CR or Rsv supplementation induced a female-specific decrease of FC between SubC-Hipp, Sub-DMLN and SubC. Moreover, Rsv supplementation lowered FC within the Hipp network and between Hipp-DMLN, Hipp-LCN and Hipp-Sens, as well as between the SubC-LCN and SubC-Sens – an effect not observed for the CR rats.

The core mechanism of CR bases itself on the reduction in energy expenditure, in which the slowing of the metabolic rate is hypothesized to improve metabolic health and extend lifespan. The selective use of energy is reflected in the reduction of biomass and thus BW. As energy for activity and basic function maintenance cannot change, limited energy resources are allocated to biomass resulting in weight loss (35). This weight loss has been reported in a plethora of preclinical (36) and clinical (37, 38) studies and is in line with our own findings, as we observe a significant decrease in BW in our male and female CR rats.

Along with the loss of biomass, energy expenditure declines until eventually the energy intake matches the new lower BW. This adaptive process allows for the maintenance of important body functions including metabolic homeostasis, breathing, heart rate, and most importantly, activity of the central nervous system (37). By this principle, RSN FC should remain mostly unaltered, as energy expenditure allocated to neuronal activity is maintained throughout CR. In a mixed-sex human cohort, whole-brain RSN FC analysis has shown to not be altered as a result of dietary intervention through a hypocaloric Mediterranean diet (39). Female-only human cohorts that have been submitted to this hypocaloric Mediterranean diet or CR, revealed a decrease of FC in regions associated with food reward (40), memory consolidation (23), self-perception and emotional functions (41). These processes are known to be modulated by hormones, with fluctuations in FC coinciding with the female menstrual cycle (42-44). This sex-related decrease in FC supports our own findings, as our results similarly show lower FC in CR females when compared to Ctrl females, between the Hipp-SubC, SubC-DMLN and within the SubC, which are networks involved with learning, memory and emotion, suggesting an underlying sex-specific mechanism of CR (45) on RSN FC. However, to our knowledge, the effects of CR on FC have not been explored in male-only cohorts. Consequently, we can’t conclusively determine if these observations are indeed sex-specific. Further exploration of the sex-specific mechanisms of CR on brain activity is therefore detrimental to understand and optimize its therapeutic potential, tailoring interventions to individual needs.

CR-mimetics are compounds that are able to increase life- and/or health-span and ameliorate age-associated diseases in model organisms, in a CR-like manner (12). Rsv as a proposed CR-mimetic has been implicated with some controversy, with conflicting reports regarding the aforementioned criteria. Rsv supplementation studies in humans do not yield overwhelmingly positive data regarding aging-related benefits (46), even though Rsv has been reported to address age- and disease-related underlying mechanisms such as systemic inflammation and oxidative stress (47, 48). Studies in model organisms using CR or Rsv on age- and obesity-related biomarkers have found small, non-existent, and even opposing effects of Rsv compared to CR mechanisms (11, 49). One of the obesity-related biomarkers in which contradicting results have been reported is BW (50). A handful of preclinical trials in rats and mice supplemented daily with Rsv (5-24 mg/kg, oral, 30-45 days), have shown unaltered (51, 52) or reduced BW (53). In this study we did not observe a significant effect of Rsv supplementation (10 mg/kg, oral) on the percentage change of BW when compared to Ctrl. However, we did observe a higher percentage increase of BW for male Rsv supplemented rats relative to their starting weight when compared to Rsv supplemented females. This difference of percentage change in BW between males and females is absent in the Ctrl and CR group. To our knowledge, there is currently no literature to support or contradict our observation of this sex-specific effect of Rsv regarding BW. Therefore, further investigation into this aspect and the effect of Rsv on BW in different sexes is warranted.

Supporting the narrative that Rsv is a CR-mimetic, we observed a female-specific decrease in the Hipp-SubC and SubC-LCN connectivity as a result of Rsv supplementation, similar to the decrease observed in female CR rats when compared to Ctrl. However, unlike CR, we observed a treatment effect of Rsv, decreasing FC within the Hipp and between the Hipp, the SubC, and other RSNs compared to Ctrl rats. Despite limited literature on the effects of Rsv on FC, published works in humans manage to directly contradict our reported lowering of FC because of Rsv supplementation. They report increased FC between the hippocampus and other regions including the right and lateral angular cortex, anterior cingulate cortex, precuneus and lateral occipital cortex, speculated to be linked to improved memory retention and attenuated hippocampal atrophy (25, 26). One possible explanation for the observed differential effects of Rsv on FC lies within the experimental design, more specifically, the dosage of Rsv. In both the mentioned human trials as well as our study, Rsv was supplemented orally daily, for the duration of a month. However, their Rsv dosage was 200 mg/kg: 20 times more compared to our daily dosage (10 mg/kg). Rsv is known to exert a hormesis dose-dependent effect (54, 55), with this effect referring to the biphasic response of a cell or organism to a compound. Often, a stimulating effect can be observed at low doses (often associated with beneficial effects), while an inhibitory effect is exerted at high doses (often toxic) (56, 57). The biphasic effect of Rsv has been shown in a plethora of (pre)clinical studies, with a multitude of them reporting differential outcomes depending on the supplied Rsv dosage (58-61). As Rsv demonstrates hormesis in various biological models, further work is required to understand its dose-relationship in context of its exerted effects, highlighting the importance of carefully considering dosage when designing studies involving Rsv supplementation and interpreting their outcomes.

The BOLD signal, used in this study as an indirect measure of neuronal activity, is based on neurovascular coupling, making it heavily dependent on changes in cerebral blood flow (CBF), cerebral blood volume (CBV) and the rate of oxygen consumption in response to changes in neuronal activity (62, 63). Unarguably, it is clear that vascular and neuronal signals both contribute to the BOLD signal (64) and while changes in BOLD are caused by neuronal activation through neurovascular coupling, they can also arise from other physiological processes that affect blood oxygenation or volume (65). CR and Rsv are both known to influence the vasculature by modulating vasodilation and CBF, through the increase of endothelial nitric oxide (NO) production (66) (67). NO plays a big role in cellular oxygen supply and demand, through regulation of the vascular tone and blood flow, as well as modulating mitochondrial oxygen consumption (68). With the increased production of endothelial NO, CR has been shown to decrease arterial blood pressure (69), improve endothelium-dependent vasodilation (70) and increase CBF (20). Similarly, Rsv supplementation leads to a more efficient endothelium-dependent vasodilation (71) and consequently, an improved CBF (72). Considering these vascular effects, it is possible they contribute to changes in the BOLD signal, which should be considered when interpreting BOLD outcomes in the context of interventions like CR and Rsv supplementation. Future work is encouraged to unravel the vascular contribution to the BOLD signal in the context of these interventions.

Sex differences in RSNs have been thoroughly investigated in humans and have shown to be confounded by environmental and sociocultural factors (73). However, sex-differences are often driven by biological factors such as hormones (e.g. estrogen) that are known to exert region-specific effects in the brain, making them potential contributors to the wide range of differences in FC observed (74, 75). Studies often show greater FC in (sub)cortical regions in females compared to males (76) and in addition, females generally show greater between-network FC whereas men have greater within-network FC (77). RsfMRI studies in rats have revealed sex-differences in brain connectivity patterns, with females exhibiting stronger hypothalamus connectivity, while males show more prominent striatum-related connectivity (78). These cumulative findings highlight the importance of taking sex into consideration as a factor when interpreting rsfMRI results in rodents, providing valuable insights into the large-scale functional organization of the brain across between sexes.

As with the majority of studies, the design of the current study is subject to limitations. First, the rsfMRI scans in this study were performed in isoflurane- and medetomidine-anesthetized rats. Anesthesia indeed affect neural activity and are likely to have an impact on FC patterns as observed in rsfMRI studies. The combination of isoflurane and medetomidine specifically is known to reduce FC in the subcortical structures such as the hippocampus and (hypo)thalamus. However, studies have shown that FC under this anaesthetics shows a very similar FC pattern as is observed in awake rats (79). Moreover, the combination of isoflurane and medetomidine has shown its own advantages, allowing visualization of interactions in cortical and sub-cortical structures, and the interaction between cortical, striatal and thalamic components (80, 81) and is therefore proposed as the most suitable anaesthetic alternative to awake imaging for rsfMRI analysis in rats (31). Secondly, rats were supplemented Rsv daily using dried apple chips to assure oral consumption. Unfortunately, most studies have shown that the oral bioavailability of resveratrol is low (<1%) (82). Consequently, this may be the reason for discrepancies between in *vivo* and *in vitro* studies regarding Rsv efficacy and mechanisms of action (83). The limited bioavailability is often accredited to several factors including poor water solubility, limited chemical stability and high metabolism (18). However, as a polyphenol, Rsv exhibits a lipophilic nature, allowing the compound to cross the blood-brain barrier and thus reach neural tissue (84), providing the speculated beneficial effects as observed in *in vivo* studies. Studies have been able to detect very low Rsv concentrations in neuronal tissue (85, 86), or report that the concentration of Rsv was too low to be detected (51), despite all establishing a wide range of beneficial effects of the compound. Thus, the data suggests that even though there is limited bioavailability of the compound, it is able to exert biological effect. It is essential to consider these limitations and explore alternative delivery methods or formulations to enhance bioavailability. Additionally, further research is needed to elucidate the precise mechanisms underlying the observed effects of Rsv to validate its therapeutic potential.

Continued research into the effects of CR and Rsv on FC using advanced neuroimaging techniques like rsfMRI holds promise for advancing our understanding of dietary interventions in the promotion of brain health. Our work provides valuable insights into the effects and comparability of short-term dietary interventions using CR and Rsv supplementation on RSNs and BW in both male and female F344 rats. With this, we established a benchmark of the sex-specific impact of CR or Rsv as a dietary intervention on spontaneous brain activity, providing an FC reference for future research of dietary effects.

## Author Contributions

Judith van Rooij: conceptualization, data curation, formal analysis, investigation, project administration, visualization, validation, writing (original draft), writing (review & editing). Monica van den Berg: visualization, conceptualization, formal analysis, project administration, software, supervision, validation, writing (review & editing). Tamara Vasilkovska: formal analysis, software, supervision, validation, writing (review & editing). Johan Van Audekerke: methodology, resources, software. Lauren Kosten: conceptualization, investigation. Daniele Bertoglio: visualization, supervision, writing (review & editing). Mohit Adhikari: visualization, methodology, resources, software, supervision, validation, writing (review & editing). Marleen Verhoye: conceptualization, project administration, resources, methodology, supervision, validation, writing (review & editing)

## Ethics approval

All experiments and handling were done in accordance with the EU legislation regulations (EU directive 2010/63/EU) and were approved by the Ethical Committee for Animal testing of the University of Antwerp (ECD # 2021-59).

