## Supplementary material for "Short-term caloric restriction or resveratrol supplementation alters large-scale brain network connectivity in male and female rats"

**Supplementary Table 1.** Brain regions and Network abbreviations

| Network | Region | Abbreviation |
|---|---|---|
| **Default mode-like network (DMLN)** | Cingulate cortex left | Cg L |
| | Cingulate cortex right | Cg R |
| | Retrosplenial cortex left | RSC L |
| | Retrosplenial cortex right | RSC R |
| | Prelimbic cortex left | Limbic L |
| | Prelimbic cortex right | Limbic R |
| | Temporal Association left | TeA L |
| | Temporal Association right | TeA R |
| **Hippocampal network (Hipp)** | Hippocampal field CA1 left | CA1 L |
| | Hippocampal field CA1 right | CA1 R |
| | Hippocampal field CA3 left | CA3 L |
| | Hippocampal field CA3 right | CA3 R |
| | Dentate gyrus left | DG L |
| | Dentate gyrus right | DG R |
| | Entorhinal cortex left | Ent L |
| | Entorhinal cortex right | Ent R |
| **Sensory network (Sens)** | Visual cortex left | VC L |
| | Visual cortex right | VC R |
| | Auditory cortex left | Aud L |
| | Auditory cortex right | Aud R |
| | Piriform cortex left | Pir L |
| | Piriform cortex right | Pir R |
| **Lateral cortical network (LCN)** | Primary somatosensory cortex left | S1 L |
| | Primary somatosensory cortex right | S1 R |
| | Secondary somatosensory cortex left | S2 L |
| | Secondary somatosensory cortex right | S2 R |
| | Motor cortex left | MC L |
| | Motor cortex right | MC R |
| | Frontal association cortex left | FrA L |
| | Frontal association cortex right | FrA R |
| | Insular cortex left | Ins L |
| | Insular cortex right | Ins R |
| **Subcortical network (SubC)** | Caudate putamen left | CPu L |
| | Caudate putamen right | CPu R |
| | Nucleus Accumbens left | NAcc L |
| | Nucleus Accumbens right | NAcc R |
| | Basal forebrain | BFB |
| | Thalamus left | Thal L |
| | Thalamus right | Thal R |
| | Medial septum left | MS L |
| | Medial septum right | MS R |
| | Hypothalamus left | Hyp L |
| | Hypothalamus right | Hyp R |

**Supplementary Table 2.** Statistical outcomes of the one-sample t-test analysis for the males of each within and between network zFC per treatment.

| Network | One-sample t-test | | | | | |
|---|---|---|---|---|---|---|
| | Males | | | | | |
| | Ctrl | | Rsv | | CR | |
| | *FC mean ± SD* | *FDR corrected p-value* | *FC mean ± SD* | *FDR corrected p-value* | *FC mean ± SD* | *FDR corrected p-value* |
| **DMLN-DMLN** | 0.497 ± 0.117 | 0.0029* | 0.379 ± 0.094 | <0.0001* | 0.527 ± 0.268 | 0.0126* |
| **DMLN-Hipp** | 0.401 ± 0.108 | 0.0029* | 0.290 ± 0.076 | <0.0001* | 0.391 ± 0.170 | 0.0079* |
| **DMLN-Sens** | 0.491 ± 0.130 | 0.0029* | 0.352 ± 0.119 | <0.0001* | 0.505 ± 0.211 | 0.0076* |
| **DMLN-LCN** | 0.269 ± 0.081 | 0.0029* | 0.187 ± 0.076 | <0.0001* | 0.301 ± 0.163 | 0.0147* |
| **DMLN-SubC** | 0.1 ± 0.062 | 0.004* | 0.134 ± 0.033 | <0.0001* | 0.199 ± 0.066 | 0.0076* |
| **Hipp-Hipp** | 0.355 ± 0.111 | 0.0029* | 0.279 ± 0.071 | <0.0001* | 0.328 ± 0.138 | 0.0076* |
| **Hipp-Sens** | 0.414 ± 0.127 | 0.0029* | 0.314 ± 0.086 | <0.0001* | 0.406 ± 0.142 | 0.0070* |
| **Hipp-LCN** | 0.265 ± 0.084 | 0.0029* | 0.207 ± 0.062 | <0.0001* | 0.295 ± 0.113 | 0.0076* |
| **Hipp-SubC** | 0.150 ± 0.059 | 0.0055* | 0.124 ± 0.022 | <0.0001* | 0.169 ± 0.053 | 0.0070* |
| **Sens-Sens** | 0.564 ± 0.190 | 0.0033* | 0.415 ± 0.120 | <0.0001* | 0.570 ± 0.178 | 0.0070* |
| **Sens-LCN** | 0.362 ± 0.106 | 0.0029* | 0.283 ± 0.072 | <0.0001* | 0.394 ± 0.160 | 0.0076* |
| **Sens-SubC** | 0.180 ± 0.061 | 0.0043* | 0.149 ± 0.031 | <0.0001* | 0.194 ± 0.062 | 0.0070* |
| **LCN-LCN** | 0.469 ± 0.065 | 0.0001* | 0.413 ± 0.043 | <0.0001* | 0.521 ± 0.212 | 0.0076* |
| **LCN-SubC** | 0.147 ± 0.043 | 0.0028* | 0.125 ± 0.035 | <0.0001* | 0.175 ± 0.061 | 0.0070* |
| **SubC-SubC** | 0.113 ± 0.037 | 0.003* | 0.093 ± 0.024 | <0.0001* | 0.136 ± 0.037 | 0.0076* |

**Supplementary Table 3.** Statistical outcomes of the one-sample t-test analysis for the females of each within and between network zFC per treatment.

| | **One-sample t-test** | | | | | |
|---|---|---|---|---|---|---|
| **Network** | **Females** | | | | | |
| | **Ctrl** | | **Rsv** | | **CR** | |
| | *FC mean ± SD* | *FDR corrected p-value* | *FC mean ± SD* | *FDR corrected p-value* | *FC mean ± SD* | *FDR corrected p-value* |
| **DMLN-DMLN** | 0.620 ± 0.149 | 0.0002* | 0.482 ± 0.174 | <0.0001* | 0.473 ± 0.134 | <0.0001* |
| **DMLN-Hipp** | 0.455 ± 0.117 | 0.0002* | 0.370 ± 0.108 | <0.0001* | 0.380 ± 0.056 | <0.0001* |
| **DMLN-Sens** | 0.565 ± 0.121 | <0.0001* | 0.464 ± 0.099 | <0.0001* | 0.456 ± 0.120 | <0.0001* |
| **DMLN-LCN** | 0.359 ± 0.040 | <0.0001* | 0.286 ± 0.079 | <0.0001* | 0.245 ± 0.093 | 0.0006* |
| **DMLN-SubC** | 0.240 ± 0.048 | <0.0001* | 0.171 ± 0.042 | <0.0001* | 0.179 ± 0.061 | 0.0002* |
| **Hipp-Hipp** | 0.413 ± 0.096 | 0.0002* | 0.336 ± 0.112 | <0.0001* | 0.371 ± 0.090 | <0.0001* |
| **Hipp-Sens** | 0.453 ± 0.106 | 0.0002* | 0.379 ± 0.078 | <0.0001* | 0.397 ± 0.060 | <0.0001* |
| **Hipp-LCN** | 0.325 ± 0.063 | <0.0001* | 0.261 ± 0.086 | <0.0001* | 0.235 ± 0.064 | <0.0001* |
| **Hipp-SubC** | 0.202 ± 0.042 | <0.0001* | 0.146 ± 0.048 | <0.0001* | 0.163 ± 0.043 | 0.0002* |
| **Sens-Sens** | 0.590 ± 0.136 | 0.0002* | 0.504 ± 0.052 | <0.0001* | 0.506 ± 0.114 | <0.0001* |
| **Sens-LCN** | 0.414 ± 0.086 | <0.0001* | 0.345 ± 0.062 | <0.0001* | 0.311 ± 0.120 | 0.0006* |
| **Sens-SubC** | 0.237 ± 0.039 | <0.0001* | 0.165 ± 0.028 | <0.0001* | 0.189 ± 0.063 | 0.0002* |
| **LCN-LCN** | 0.579 ± 0.125 | <0.0001* | 0.445 ± 0.124 | <0.0001* | 0.380 ± 0.128 | 0.0004* |
| **LCN-SubC** | 0.208 ± 0.036 | <0.0001* | 0.139 ± 0.040 | <0.0001* | 0.142 ± 0.062 | 0.0002* |
| **SubC-SubC** | 0.160 ± 0.029 | <0.0001* | 0.110 ± 0.029 | <0.0001* | 0.130 ± 0.042 | 0.0010* |

**Supplementary Table 4.** Statistical outcomes Network-based FC analysis for each within and between network zFC. In case of a non-significant sex*treatment interaction. a two-way ANOVA was performed with main effects (sex. treatment) only.

| | Two-way ANOVA | | | | | | | | | |
|---|---|---|---|---|---|---|---|---|---|---|
| **Network** | | **DF** | **Sum of squares** | **F ratio** | **P-value** | | **DF** | **Sum of squares** | **F ratio** | **P-value** |
| **DMLN-DMLN** | *Sex*treatment* | 2 | 0.080 | 1.658 | 0.2074 | *Sex* | 1 | 0.022 | 0.866 | 0.3588 |
| | | | | | | *Treatment* | 2 | 0.107 | 2.110 | 0.1377 |
| **DMLN-Hipp** | *Sex*treatment* | 2 | 0.023 | 1.436 | 0.2554 | *Sex* | 1 | 0.169 | 2.028 | 1.650 |
| | | | | | | *Treatment* | 2 | 0.119 | 7.147 | 0.0030* |
| **DMLN-Sens** | *Sex*treatment* | 2 | 0.043 | 1.225 | 0.3075 | *Sex* | 1 | 0.022 | 1.228 | 0.2757 |
| | | | | | | *Treatment* | 2 | 0.090 | 2.508 | 0.0968 |
| **DMLN-LCN** | *Sex*treatment* | 2 | 0.060 | 3.862 | 0.0326* | | | | | |
| **DMLN-SubC** | *Sex*treatment* | 2 | 0.020 | 5.243 | 0.0116* | | | | | |
| **Hipp-Hipp** | *Sex*treatment* | 2 | 0.008 | 0.671 | 0.5194 | *Sex* | 1 | 0.018 | 3.032 | 0.0922 |
| | | | | | | *Treatment* | 2 | 0.084 | 6.869 | 0.0036* |
| **Hipp-Sens** | *Sex*treatment* | 2 | 0.011 | 0.776 | 0.4698 | *Sex* | 1 | 0.014 | 1.972 | 0.1704 |
| | | | | | | *Treatment* | 2 | 0.094 | 6.214 | 0.0055* |
| **Hipp-LCN** | *Sex*treatment* | 2 | 0.021 | 2.075 | 0.1432 | *Sex* | 1 | 0.000 | 0.087 | 0.7694 |
| | | | | | | *Treatment* | 2 | 0.037 | 3.360 | 0.0474* |
| **Hipp-SubC** | *Sex*treatment* | 2 | 0.011 | 4.242 | 0.0250* | | | | | |
| **Sens-Sens** | *Sex*treatment* | 2 | 0.036 | 1.042 | 0.3647 | *Sex* | 1 | 0.004 | 0.240 | 0.6271 |
| | | | | | | *Treatment* | 2 | 0.088 | 2.481 | 0.0991 |
| **Sens-LCN** | *Sex*treatment* | 2 | 0.033 | 1.593 | 0.2199 | *Sex* | 1 | 0.000 | 0.016 | 0.8972 |
| | | | | | | *Treatment* | 2 | 0.044 | 2.029 | 0.1480 |
| **Sens-SubC** | *Sex*treatment* | 2 | 0.008 | 1.870 | 0.1728 | *Sex* | 1 | 0.005 | 2.303 | 0.1395 |
| | | | | | | *Treatment* | 2 | 0.026 | 5.764 | 0.0076* |
| **LCN-LCN** | *Sex*treatment* | 2 | 0.094 | 3.044 | 0.0621 | *Sex* | 1 | < 0.0001 | 0.000 | 0.9955 |
| | | | | | | *Treatment* | 2 | 0.071 | 2.048 | 0.1451 |
| **LCN-SubC** | *Sex*treatment* | 2 | 0.012 | 3.079 | 0.0608 | *Sex* | 1 | 0.000 | 0.257 | 0.6152 |
| | | | | | | *Treatment* | 2 | 0.017 | 3.768 | 0.0339* |
| **SubC-SubC** | *Sex*treatment* | 2 | 0.007 | 4.393 | 0.0215* | | | | | |

**Supplementary Table 5.** Statistical post-hoc (Tukey HSD) outcomes of network-based zFC analysis for each within and between network zFC. showing a significant treatment main effect.

| | **Pairwise comparisons Tukey HSD (treatment)** | | | | | | |
|---|---|---|---|---|---|---|---|
| **Network** | **Treatment** | **Difference** | **Std Error** | **t-value** | **P-value** | **Lower 95%** | **Upper 95%** |
| **DMLN-Hipp** | *CR-Ctrl* | -0.076 | 0.039 | -1.91 | 0.1545 | -0.174 | 0.022 |
| | *CR-Rsv* | 0.071 | 0.038 | 1.88 | 0.1637 | -0.022 | 0.165 |
| | *Ctrl-Rsv* | 0.147 | 0.039 | 3.78 | 0.0020* | 0.051 | 0.244 |
| **DMLN-SubC** | *CR-Ctrl* | -0.035 | 0.024 | -1.47 | 0.3175 | -0.095 | 0.023 |
| | *CR-Rsv* | 0.032 | 0.023 | 1.36 | 0.3722 | -0.026 | 0.090 |
| | *Ctrl-Rsv* | 0.067 | 0.024 | 2.81 | 0.0226* | 0.008 | 0.127 |
| **Hipp-Hipp** | *CR-Ctrl* | -0.068 | 0.034 | -2.01 | 0.1289 | -0.153 | 0.015 |
| | *CR-Rsv* | 0.055 | 0.032 | 1.70 | 0.2222 | -0.025 | 0.136 |
| | *Ctrl-Rsv* | 0.124 | 0.033 | 3.71 | 0.0025* | 0.041 | 0.207 |
| **Hipp-Sens** | *CR-Ctrl* | -0.054 | 0.037 | -1.45 | 0.3269 | -0.146 | 0.037 |
| | *CR-Rsv* | 0.075 | 0.035 | 2.12 | 0.1023 | -0.012 | 0.163 |
| | *Ctrl-Rsv* | 0.129 | 0.037 | 3.49 | 0.0043* | 0.038 | 0.221 |
| **Hipp-LCN** | *CR-Ctrl* | -0.037 | 0.031 | -1.21 | 0.4569 | -0.114 | 0.038 |
| | *CR-Rsv* | 0.041 | 0.030 | 1.38 | 0.3608 | -0.032 | 0.115 |
| | *Ctrl-Rsv* | 0.079 | 0.030 | 2.59 | 0.0375* | 0.003 | 0.154 |
| **Hipp-SubC** | CR-Ctrl | -0.036 | 0.018 | -2.02 | 0.1271 | -0.081 | 0.008 |
| | CR-Rsv | 0.030 | 0.017 | 1.81 | 0.1848 | -0.011 | 0.072 |
| | Ctrl-Rsv | 0.067 | 0.017 | 3.86 | 0.0017* | 0.024 | 0.110 |
| **LCN-SubC** | CR-Ctrl | -0.025 | 0.020 | -1.24 | 0.4404 | -0.074 | 0.024 |
| | CR-Rsv | 0.029 | 0.019 | 1.51 | 0.3014 | -0.018 | 0.077 |
| | Ctrl-Rsv | 0.054 | 0.019 | 2.73 | 0.0266* | 0.005 | 0.103 |
| **Sens-SubC** | CR-Ctrl | -0.026 | 0.020 | -1.29 | 0.4131 | -0.076 | 0.024 |
| | CR-Rsv | 0.041 | 0.019 | 2.14 | 0.0986 | -0.006 | 0.089 |
| | Ctrl-Rsv | 0.068 | 0.020 | 3.34 | 0.0062* | 0.017 | 0.118 |

**Supplementary Table 6.** Statistical post-hoc (Student t-test) outcomes of network-based zFC analysis for each within and between network zFC. showing a significant sex*treatment interaction effect.

| | **Pairwise comparisons Student T-test (sex*treatment)** | | | | | | | |
|---|---|---|---|---|---|---|---|---|
| **Network** | **Treatment** | **Difference** | **Std Error** | **t value** | **P-value** | **FDR correction** | **Lower 95%** | **Upper 95%** |
| **DMLN-LCN** | Female CR - Female Ctrl | -0.114 | 0.049 | -2.33 | 0.0272* | 0.0816 | -0.215 | -0.013 |
| | Female CR – Female Rsv | -0.041 | 0.047 | -0.88 | 0.3858 | 0.3858 | -0.138 | 0.055 |
| | Female CR – Male CR | -0.056 | 0.051 | -1.08 | 0.2878 | 0.3263 | -0.162 | 0.049 |
| | Female Ctrl – Female Rsv | 0.072 | 0.049 | 1.48 | 0.1497 | 0.2694 | -0.027 | 0.173 |
| | Female Ctrl – Male Ctrl | 0.122 | 0.057 | 2.14 | 0.0407* | 0.0916 | 0.005 | 0.239 |
| | Female Rsv – Male Rsv | 0.120 | 0.049 | 2.45 | 0.0206* | 0.0816 | 0.019 | 0.221 |
| | Male CR – Male Ctrl | 0.064 | 0.059 | 1.08 | 0.2900 | 0.3263 | -0.057 | 0.185 |
| | Male CR – Male Rsv | 0.135 | 0.053 | 2.52 | 0.0175* | 0.0816 | 0.025 | 0.244 |
| | Male Ctrl – Male Rsv | 0.071 | 0.057 | 1.24 | 0.2236 | 0.3263 | -0.045 | 0.187 |
| **Hipp-SubC** | Female CR - Female Ctrl | -0.069 | 0.021 | -3.17 | 0.0038* | 0.019* | -0.114 | -0.024 |
| | Female CR – Female Rsv | 0.017 | 0.020 | 0.83 | 0.4133 | 0.4769 | -0.025 | 0.060 |
| | Female CR – Male CR | -0.020 | 0.021 | -0.94 | 0.3533 | 0.4769 | -0.065 | 0.024 |
| | Female Ctrl – Female Rsv | 0.087 | 0.021 | 3.96 | 0.0005* | 0.0038* | 0.041 | 0.132 |
| | Female Ctrl – Male Ctrl | 0.068 | 0.022 | 2.97 | 0.0061* | 0.0229* | 0.021 | 0.115 |
| | Female Rsv – Male Rsv | 0.000 | 0.020 | 0.04 | 0.9667 | 0.9667 | -0.042 | 0.043 |
| | Male CR – Male Ctrl | 0.019 | 0.022 | 0.85 | 0.4051 | 0.4769 | -0.027 | 0.066 |
| | Male CR – Male Rsv | 0.039 | 0.021 | 1.78 | 0.0869 | 0.201 | -0.006 | 0.084 |
| | Male Ctrl – Male Rsv | 0.019 | 0.021 | 0.89 | 0.3795 | 0.4769 | -0.025 | 0.064 |
| **LCN-SubC** | Female CR - Female Ctrl | -0.089 | 0.029 | -3.09 | 0.0043* | 0.0193* | -0.148 | -0.030 |
| | Female CR – Female Rsv | 0.012 | 0.029 | 0.42 | 0.6769 | 0.7615 | -0.047 | 0.071 |
| | Female CR – Male CR | -0.025 | 0.030 | -0.84 | 0.4070 | 0.6663 | -0.088 | 0.036 |
| | Female Ctrl – Female Rsv | 0.101 | 0.030 | 3.38 | 0.0020* | 0.0180* | 0.040 | 0.163 |
| | Female Ctrl – Male Ctrl | 0.089 | 0.031 | 2.84 | 0.0081* | 0.0243* | 0.025 | 0.154 |
| | Female Rsv – Male Rsv | 0.001 | 0.029 | 0.03 | 0.9798 | 0.9798 | -0.058 | 0.060 |
| | Male CR – Male Ctrl | 0.025 | 0.033 | 0.78 | 0.4442 | 0.6663 | -0.041 | 0.092 |
| | Male CR – Male Rsv | 0.038 | 0.030 | 1.27 | 0.2156 | 0.4851 | -0.023 | 0.101 |
| | Male Ctrl – Male Rsv | 0.013 | 0.030 | 0.43 | 0.6720 | 0.7615 | -0.049 | 0.075 |
| **SubC** | Female CR - Female Ctrl | -0.052 | 0.017 | -2.96 | 0.006* | 0.0225* | -0.089 | -0.016 |
| | Female CR – Female Rsv | 0.001 | 0.016 | 0.1 | 0.9201 | 0.9201 | -0.031 | 0.035 |
| | Female CR – Male CR | -0.018 | 0.017 | -1.06 | 0.2984 | 0.373 | -0.055 | 0.017 |

| | | | | | | | | |
|---|---|---|---|---|---|---|---|---|
| | Female Ctrl – Female Rsv | 0.054 | 0.017 | 3.16 | 0.0037* | 0.0225* | 0.019 | 0.089 |
| | Female Ctrl – Male Ctrl | 0.057 | 0.018 | 3.07 | 0.0046* | 0.0225* | 0.019 | 0.095 |
| | Female Rsv – Male Rsv | 0.022 | 0.015 | 1.45 | 0.1587 | 0.2976 | -0.009 | 0.054 |
| | Male CR – Male Ctrl | 0.023 | 0.018 | 1.25 | 0.2213 | 0.3476 | -0.014 | 0.061 |
| | Male CR – Male Rsv | 0.043 | 0.017 | 2.51 | 0.0178* | 0.0534 | 0.008 | 0.078 |
| | Male Ctrl – Male Rsv | 0.020 | 0.017 | 1.16 | 0.2549 | 0.3476 | -0.015 | 0.055 |
| **DMLN-SubC** | Female CR - Female Ctrl | -0.097 | 0.026 | -3.65 | 0.0011* | 0.0083* | -0.151 | -0.042 |
| | Female CR – Female Rsv | -0.023 | 0.025 | -0.91 | 0.3691 | 0.5033 | -0.075 | 0.028 |
| | Female CR – Male CR | -0.038 | 0.026 | -1.45 | 0.1593 | 0.2655 | -0.092 | 0.016 |
| | Female Ctrl – Female Rsv | 0.073 | 0.026 | 2.78 | 0.0097* | 0.0364* | 0.019 | 0.128 |
| | Female Ctrl – Male Ctrl | 0.079 | 0.027 | 2.86 | 0.0079* | 0.0364* | 0.022 | 0.136 |
| | Female Rsv – Male Rsv | 0.050 | 0.024 | 2.06 | 0.0491* | 0.1052 | 0.002 | 0.100 |
| | Male CR – Male Ctrl | 0.020 | 0.027 | 0.75 | 0.4573 | 0.5716 | -0.035 | 0.077 |
| | Male CR – Male Rsv | 0.065 | 0.025 | 2.55 | 0.0165* | 0.0495* | 0.012 | 0.118 |
| | Male Ctrl – Male Rsv | 0.044 | 0.025 | 1.74 | 0.0936 | 0.1755 | -0.008 | 0.097 |

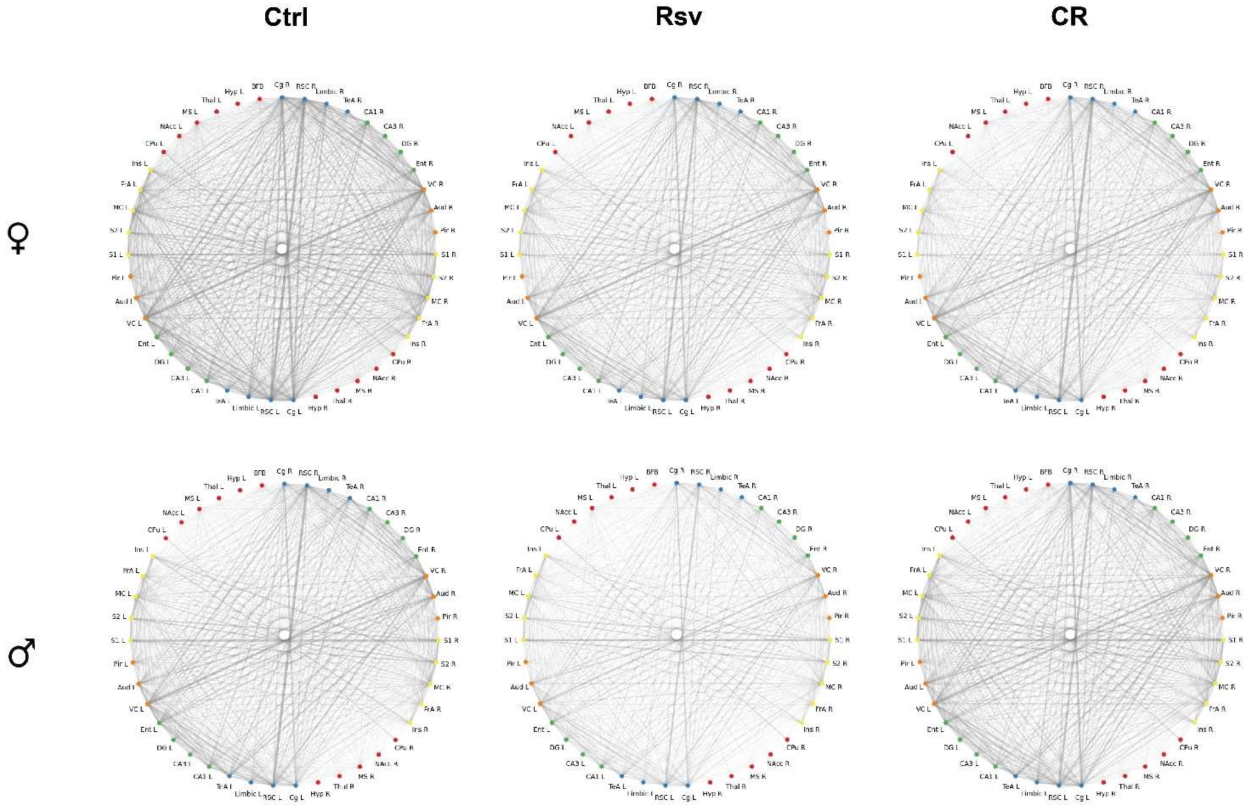


**Supplementary Figure 1.** FC in male and female Ctrl, Rsv supplemented and CR rats. Chord plots displaying the mean FC between ROI pairs for males and females per treatment group (Ctrl = control, CR = caloric restriction, Rsv = Resveratrol. The width and transparency of the edges indicate the strength of FC. Broad and dark grey edges indicate high FC and narrow light grey edges indicate low FC. Non-significant connections ($p > 0.05$, one-sample t-test, FDR corrected per group) are not shown. The colours of the nodes indicate the corresponding RSNs of the ROIs: DMLN (default mode-like network = blue), Hipp (hippocampal network = green), Sens (sensory network= orange), LCN (lateral cortical network = yellow), SubC (subcortical network = red).
